# PEPsRNA: A computational resource of experimentally tested peptide based siRNA delivery

**DOI:** 10.64898/2026.02.12.705561

**Authors:** Showkat Ahmad Dar, Manoj Kumar

**Author notes:** Correspondence (Temporary).

## Abstract

In siRNA-based applications, cellular delivery remains one of the main hurdles. Many formulations are tested for the same and peptides came up as one of the optimal options. The latter have various advantages like natural biological presence, high specificity, and natural metabolism etc. siRNA in conjugation with peptides have exhibited enhanced mRNA silencing. Peptides aid siRNAs in condensation to smaller volumes, enhance nuclease protection, increase half-life, promote cell specific binding as well as endosomal escape and release in cytosol. Despite its prime importance, no resource is available for the peptide-based delivery of siRNAs, therefore to fill the gap we developed PEPsRNA web server. It includes 2266 entries of 270 different kinds of peptides, 106 different types of siRNAs and shRNAs along with more than 80 conjugate molecules targeting 55 different genes, experimentally tested for the delivery of the siRNAs. To provide the detailed insights of the procedure, we have incorporated analysis of the peptides (e.g. secondary structure, amino acid composition, polarity, hydrophobicity etc.), siRNAs (e.g. secondary structures with minimum free energies etc.) and associated conjugate molecules (e.g. structure, SMILES, Inchl). We have derived these values using various other tools and resources to make the web server comprehensive. We further compared various physicochemical properties with the efficacy of the peptide based on the target gene silencing, but these properties do not shown any distinct conclusive relationship. The data is available for browsing, searching and downloading freely on the web server with URL: http://bioinfo.imtech.res.in/manojk/pepsirna.

**Highlights:**

➣ PEPsRNA is the first database of experimentally tested peptides for siRNA delivery
➣ It comprised of 2266 entries with 270 peptides and about 80 conjugate molecules
➣ Analysis of peptides, siRNAs and details of conjugate molecules are provided
➣ Browse, search and various tools are incorporated for data retrieval and usage

## Introduction

With the discovery of RNA interference (RNAi) phenomena by “Andrew Fire” and coworkers, it became the center of attention for silencing the target genes [1]. Double stranded, 21 nucleotide long RNA molecules called small interfering RNA (siRNA) binds to the “RNA-induced silencing complex” (RISC) in cytoplasm and recognizes its target mRNA in sequence specific manner that leads to the degradation of latter [2]. These siRNAs are used extensively in research for single gene wise or genome wide level silencing [3]. The RNAi technology is also extensively investigated as potential therapeutic for abnormal conditions ranging from deadly virus-causing pathologies, metastasis and genetic disorders [4, 5]. Various experiments have proved its ability to treat these abnormalities at least in labs and many siRNA-based molecules are in clinical trials pipeline [6, 7]. Furthermore, with the engineering of chemical modifications or abasic siRNAs, advancement in siRNA based therapeutics has progressed which is evident with the FDA approval of the Patisiran (ONPATTRO™) [8-10]. This approval has developed new impetus in the field of siRNA based therapeutics development and research. However, there are certain limitations for in vivo use of the siRNAs and delivery is one of the main hurdles [8, 11]. Some of the main barriers that are encountered for siRNA delivery include (a) the condensation of the siRNAs into compact units to lessen the unnecessary interaction and protection from the nucleases, (b) cell specific delivery i.e. less off targets and use of minimal inhibitory concentration, (c) cell specific uptake, (d) endosomal escape and (e) siRNA release in cytoplasm as active molecule [12, 13]. Various types of natural and synthetic complexes are attempted for the development of siRNA delivery vehicle e.g. peptides, exosomes, chitosan, dendrimers, liposomes, aptamers, polyethylene glycol, polyethyleneimine, gold particles etc. [14, 15].

Peptides are promising delivery vectors for siRNAs either as peptide-siRNA complex or in combination with other molecules to form nano-formulations [16]. The siRNAs are negatively charged molecules and therefore will repel each other (and cell membranes), which will in turn lead to an increase in the siRNA-particle size. Positively charged amino acids like cationic peptides (rich in Arginine), including poly(L-lysine) (PLL), protamine etc. have been used to mitigate this limitation [17]. For example, protamine and similar recombinant peptides condense the siRNAs in size and also significantly increased its half-life and cellular delivery [18]. Cell membrane penetration and intracellular delivery of siRNA is achieved by the active cell penetrating peptides [19]. But in the systemic delivery, the technique works on the basis of concentration gradient and has minimal effect on the distant cells or intratumoral cells [20]. This problem is solved by the receptor specific peptide ligands that bind to the specific receptors and result in the siRNA internalization by corresponding cells only [20]. The combination of such peptides with the lipid molecule (like myristic acid) can even cross the blood brain barrier [21]. One such example is the use of “cell-penetrating peptide (transportan)”, “receptor-targeting peptide (transferrin)” and “myristic acid” that was efficiently able to deliver active siRNA in the “brain cells” with resistance against serum ribonucleases [21]. Receptor specific ligand peptides are also used in siRNA-liposome complex on their surfaces to promote the cell specific delivery [22]. Subsequently, inside the cells endosomal localization of the siRNA (or its complex) occurs that subsequently degrade the siRNA molecule hence rendering it ineffective [23]. This phenomenon is avoided by endosome-lysing peptides or proton-gradient, generated by histidine rich peptides that lyse the endosome placing the siRNA in the cytoplasm [24]. Also, the peptide-siRNAs are covalently linked with some protease recognizing linker that is cleaved only when it reaches inside the cell e.g. Cathepsin-D (rich in cancer cell endosomes) recognizing peptide is used as a linker of siRNA and peptide [25].

The process of siRNA delivery is a multistep process and requires likewise approach for efficient delivery of the RNA molecule. As seen above, peptides can play such roles at every step in the targeted delivery of the siRNA based silencing. We looked in the literature for resources on delivery of siRNA (including peptides for their delivery) or relevant molecules but could not find any. However, number of resource regarding RNAi as well as peptides (disjointedly) are present like VIRsiRNAdb (viral siRNA database)[26], HuSiDa (Human specific siRNAs)[27], siRNAdb (database of siRNA sequences) for small RNAs [28], HTS-DB (high-throughput data of siRNA screening projects[29], PVsiRNAdb (database for plant exclusive virus-derived small interfering RNAs)[30]. Likewise, CPPsite-2.0 (cell-penetrating peptides)[31], CancerPPD (anticancer peptides and proteins)” [32], “APD3 (Antimicrobial Peptide Database)[33], AHTPDB (antihypertensive peptides)[34], SATPdb (structurally annotated therapeutic peptides) [31] for peptides. Also, none of the resource examines both siRNAs and peptides (or other associated molecules) collectively. Therefore, after confirming the importance of the peptide molecules in the medical formulation and their potential to develop the delivery vehicle for siRNAs, we tried to fill the gap and developed “PEPsRNA” web server.

## Results

### Data browsing and searching

We developed PEPsRNA web server, a first resource that incorporated the data of both siRNAs and the peptides (including the conjugate molecules) that are used specifically for delivery. The overall organization of the PEPsRNA web server is shown in **Figure 1**. To explore the PEPsRNA data, start from “Browse” option within which main fields are listed including peptide, target gene, conjugate molecule, experiment used and the study (title) with reference PubMed Id (PMID). Further in each of these categories, a unique list of corresponding entries is provided e.g. in “peptide-option,” list of peptides is presented. These peptides can further be clicked into the sub-list of entries in which that particular peptide exists. Further, clicking on the database-id (e.g. PEPsiRNA1575) of a particular entry, the page displaying the general information about the same entry.

**Figure 1.**
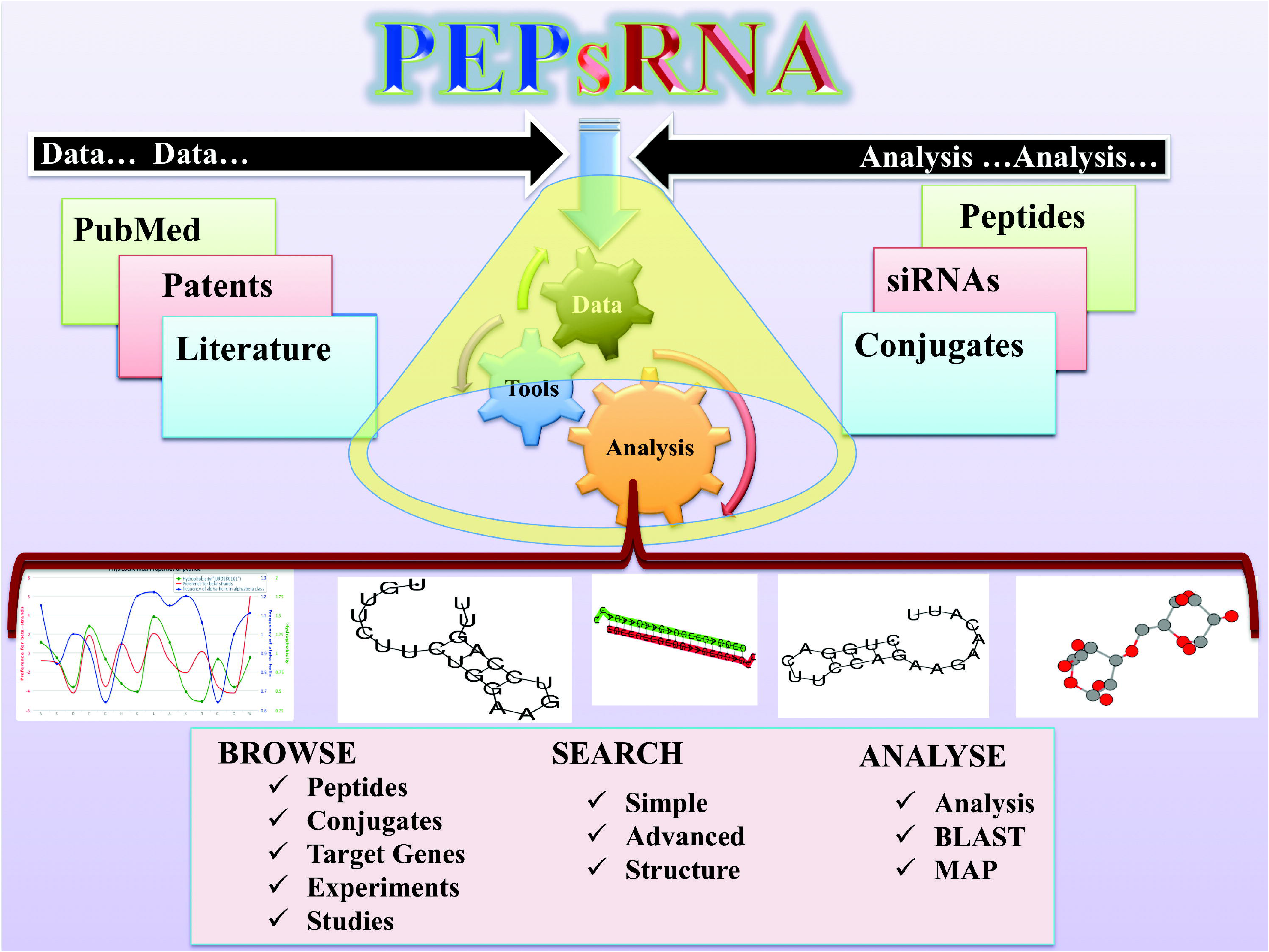
The organizational chart of PEPsRNA web server. Data extraction and analysis was performed with all the details of siRNA, peptide and conjugate provided in the web server with Browse, Search and Analysis options.

Similarly, for data searching, three options are provided under “Search” option. The first one is the “Simple search” which involves many categories for “string (word)” search. The various categories listed in this option are Reference, Peptide Sequence, Peptide name, Target gene, Efficacy of peptide, Gene expression, Conjugate molecule, SMILE, Experiment, Cell or Organism used, Source, Author. The next search option is “Advanced search” in which user can explore the combinations of properties like “Peptide” and “Conjugate molecule” for searching the database. Finally “Structure based search” is provided specifically for conjugate molecules, in which the user can draw the structure and search it in the database. Next, alternatives for customized BLAST (Basic local alignment search tool) based search of peptides, siRNA, peptide secondary structures or the analysis of the peptide etc. are provided under the “Tools” web page on web server. Likewise, “Map” option is provided for siRNA and peptides that will check input sequences provided by the user for all the sequences in the database and fetch them if present in the database. The “Help page” is provided to ease the user to become familiar with the web server.

### PEPsRNA database statistics

We analyzed the siRNA delivery peptides for their lengths and they fall in the range of 5 to 40 amino acids however, the long peptides up to stretch of 153 amino acids were also found **Figure 2 (a**). Also, these peptides fall into two categories in relation to linking with the siRNAs molecules namely covalent or non-covalent (electrostatic) linking **Figure 2 (b**). The statistical analysis of the PEPsRNA database shows that most of the experiments were performed in vitro systems and about 15 percent of tests done in live models (mainly mice). The HeLa cells followed by A549 cells were the most common cell lines used for testing these peptides among 92 different cell lines. **Supplementary Figure S4** shows the diagrammatic representation of the experimental type and cell line dispersal used for testing the peptide based siRNA delivery.

**Figure 2.**
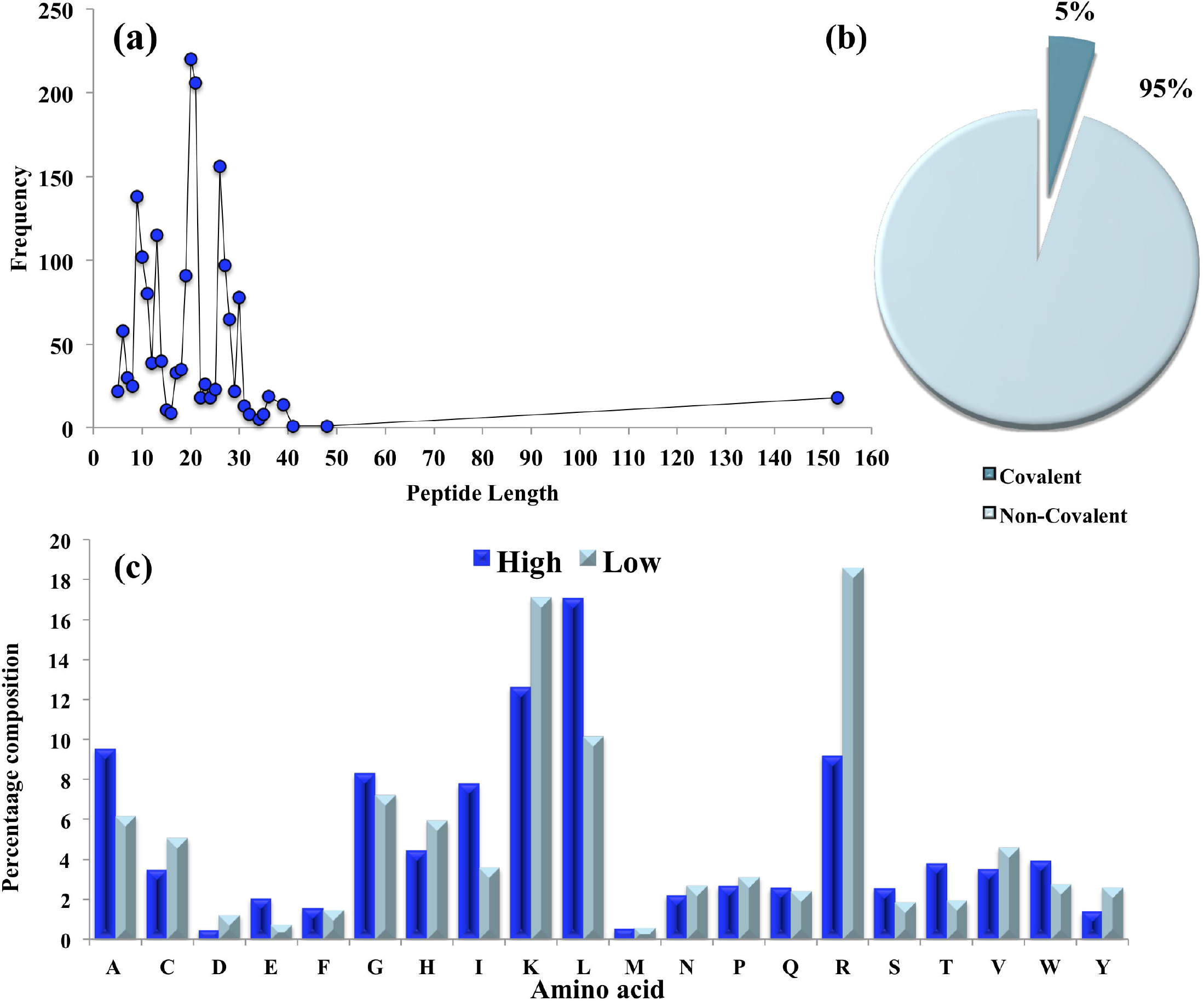
Peptide frequencies and their interaction with siRNAs. (a) Lengths wise frequency of the peptides used for delivery of the siRNAs. (b) Linking information between the peptides and the siRNAs. (c) Amino acid frequency analysis of high and low efficacy delivery peptides (80 sequences of each category).

The target genes to be silenced by the peptide-based siRNAs were 55 and luciferase was the most common gene used as shown in **Supplementary Figure S5** (**a**). The distribution of commonly used conjugate molecules (other than peptides) is shown in **Supplementary Figure S5** (**b**), and Polyethylene glycol (PEG) tops the number among 75 other such molecules.

### Analysis of the delivery peptides

For the analysis of main physicochemical properties of peptides, we plotted them against the delivery efficacies of the respective peptides that were deduced from the knockdown of the target gene. The scatter plot matrix of various properties like Efficacy, Length, Cell penetrating peptides score, isoelectric point, Charge, Hydrophilicity and Hydrophobicity (**Figure 3**). The individual scatter plots are shown in supplementary material (**Supplementary Figures S6, S7**). Additionally, from amino acid composition analysis of the 80 sequences from very high and very low efficacy peptides, we found that Leucine (L), Lysine (K), Alanine (A), Arginine (R) and Glycine (G) amino acids predominated both kinds of peptides (**Figure 2(c)**).

**Figure 3.**
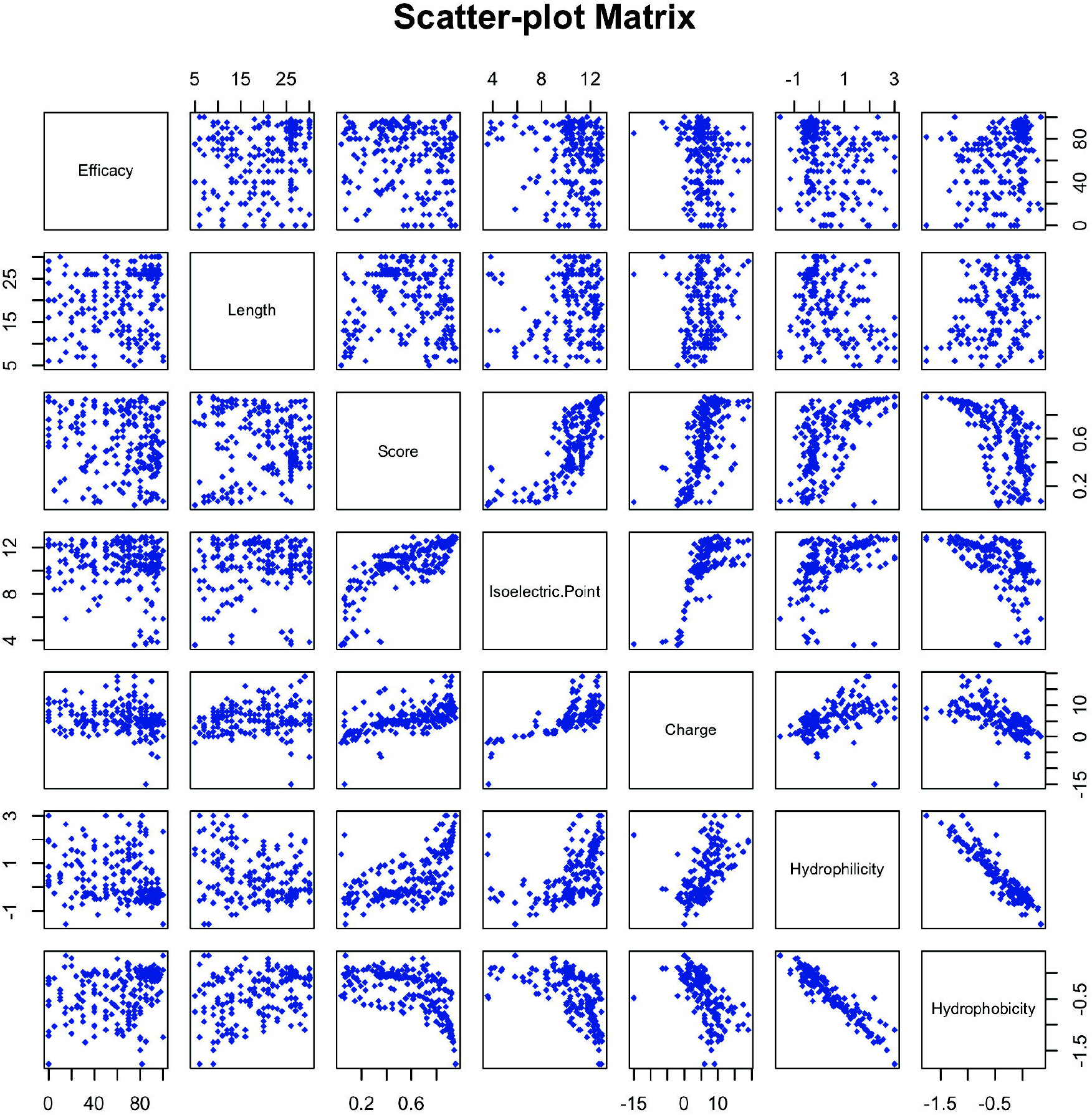
The scatter-plot matrix of various physicochemical properties of the peptides with each other and experimental delivery efficacy (percentage). The various inputs are (i) Efficacy, which represents experimental efficacy as derived from research papers for delivery of siRNA and silencing of the target gene, (ii) Length of the peptides, (iii) Score for predicted cell penetrating peptide probability score, (iv) Isoelectric point, (v) Charge (vi) Hydrophilicity and (vii) Hydrophobicity of corresponding peptides.

However, Leucine, Arginine, Glycine and Iso-leucine predominated in high efficacy peptides while Arginine, Lysine, Histidine, Valine, and Cysteine amino acids abounded in low efficacy peptides.

## Discussion

The PEPsRNA web server consists of 2266 entries with more than 270 unique peptides and more than 75 non-peptide conjugate molecules experimentally tested for delivery of the siRNAs. The peptides in the database not only include cell-penetrating peptides (CPP) but also cell specific binding as well as other peptides that will help the release of siRNA inside the cytoplasm [35-37]. Also, the combination of peptides with other non-peptide molecules enhances their efficiency in the delivery of siRNAs [38]. Therefore, we have considered all these factors to create repository for peptide based siRNA delivery. The broad fields of the data involved are (I) peptides, (II) siRNAs and (III) non-peptide conjugate molecules besides, other experimental information like cell lines, experiment used and inference in terms of peptide effectiveness as delivery vehicle (**Supplementary Figure S1**). For each main category and each entry, we have provided detailed analysis of their properties. For example, for each individual peptide, we have analyzed its secondary structure, composition, polarity, hydrophilicity as well as hydrophobicity, preference to the alpha helix, beta strand, amino acid surface exposure, disorder promoting percentage, flexibility etc. (**Supplementary Figure S2-a**). Similarly, we have provided the analysis in pictorial and numerical values for the double stranded siRNAs as well as for each sense and antisense strands. This includes the dot plots, secondary structures (with minimum free energy kcal/mol), line notations etc. (**Supplementary Figure S2-b**). The peptides are tested with wide range of other molecules ranging from silica or gold nanoparticles, chitosan to chemical molecules, lipids to liposomes, polyplex to polyurethanes etc. Each of these molecules is well documented with its chemical structure, SMILES, Inchl and links to PubChem (**Supplementary Figure S3**). There are various options provided in the web server to make it easy for the user to explore the data.

From the statistical point of view, mainly in vitro experiments dominated the research area and easily detectable targets like luciferase or fluorescent proteins were top most genes used in peptide based siRNA knockdown. The reason could be the easy detection of end product of these genes and their testing in cell lines rather than living systems. The peptides range from the length of 5 amino acids to about 150 amino acids, but majority of the peptides lie in between 8 to 35 amino acid length (**Figure 2-a**). The peptides were mainly free and not linked via any bonding to the siRNA except for very few siRNAs. This might be because the covalent connection may hinder siRNA release in the cytoplasm and it needs additional resources to be executed for linking and unlinking the two (peptide-siRNA) before and after their delivery into the cell.

For further analyses of the peptides, we mapped the matrix scatterplot of various physicochemical properties against each other with primary property as their experimental efficacies (in terms of target mRNA silencing), ranging form 0 to 100 percent as derived from research papers (**Figure 3, Supplementary Figures S6, S7**). The plot of predicted CPP score (defined as “Score” in figure) versus efficacy depicted that not all good efficacy peptides are CPP as a good chunk of non-CPP peptides performed better. The reason for this is that some peptides were also directed to receptors and are internalized via receptor mediated cell entry of cargo [36, 37]. Similarly, The length (limited to 30 amino acids) versus efficacy chart also showed the same pattern of not much relevance between the two. From the isoelectric point (pI) point of view, all the peptides were mainly in the range of 8 to 10 (pH) and were mainly positively charged ranging from 2 to 10. Next, the inclination of the peptides was more towards hydrophilicity (**Supplementary Figure S7**), which may be due to the presence of the arginine amino acid. The positive charge of the arginine, lysine etc. is required for neutralizing the negative charge of siRNA and for interaction with the cell membrane. Lastly, from the percentage composition and motif analysis of the peptides, we could not derive any significant compositional difference in high and low efficacy peptides (80 each). The positively charged amino acids dominated the composition in both types of peptides (**Figure 2 (c)**). However, the low efficacy peptides showed more prevalence of Arginine, Lysine and Histidine (positively charged amino acids). While, in high efficiency category Leucine, Alanine, Glycine, Isoleucine, Tryptophan (Hydrophilic side chain amino acids) and Threonine (uncharged polar side chain) dominated. Overall, the pattern of amino acid percentage composition in both types showed a similar trend of occurrence i.e. the amino acids which are more in high efficacy peptides are also more in low efficacy peptides and vice versa.

## Conclusion

Delivery remains the most important aspects in the development of siRNA-based therapeutics and peptide provide one of the best solutions for it. The number of peptide-based products is increasing in pharmaceutical market as delivery vehicles or themselves as therapeutic agents. With this in mind, we developed PEPsRNA web server, the first computational resource for siRNA delivery focused on the peptides as carrier vehicles. Detailed experimental data from research articles as well as patents for peptides, siRNAs and non-peptide conjugate molecules are analyzed and provided. We expect this web server to be of great importance to researchers working in the fields of RNAi, its delivery, peptides, siRNA based therapeutics development etc. The resource is available online freely for scientific and educational use on the URL: http://bioinfo.imtech.res.in/manojk/pepsirna.

## Materials and Methods

### Acquisition, curation and association of the data

We acquired the main data from the research articles and patents after developing the search query using advanced options in the PubMed, Patent lens (https://www.lens.org) and Google patents (https://patents.google.com) [39]. We developed the query related to keywords of “peptide” and “siRNA” as below:

“(((peptides OR peptide*))) AND (((((((RNA silencing) OR RNA interference) OR RNAi) OR shRNA) OR short interfering RNA) OR small interfering RNA) OR siRNA*)”

We got around “66500” number of articles from PubMed till start of 2019. From these vast numbers of articles, we screened the relevant articles excluding the reviews, Non-English articles etc. Most of the articles were related to siRNA based silencing and with non-relevant information regarding the peptide based delivery. Our search terms are broad in nature so that we do not miss any relevant information. We manually curated the documents and at last extracted data from “104” research articles and “5” patents. The data includes 2266 number of entries with information on peptides, siRNA sequences, non-peptide conjugate molecules used in delivery, experimental conditions and other related information. The database includes 270 unique peptides and we derived their secondary structure using the “CFSSP: Chou and Fasman Secondary Structure Prediction server (http://www.biogem.org/tool/chou-fasman/index.php)”. Besides, other properties like the amino acid composition, polarity, hydrophobicity, surface exposure, and flexibility of amino acids with pictorial representations are provided [40]. Next, we mined the chemical information of the non-peptide conjugate molecules involved, which were found to be around 75 in number. We collected their information which include their chemical structure, simplified molecular-input line-entry system (SMILES), International Chemical Identifier (InChl) and link to external repository-PubChem [8]. Similarly, the number of unique siRNAs (and shRNAs) was around 106, for which we used RNAfold package to analyze the secondary structure, thermodynamic properties etc. of each strand and double stranded siRNA molecule as well [39]. Furthermore, the unique peptides (size limit 5 to 30 amino acids) were analyzed for their cell-penetrating efficacy using the online prediction server “CPPpred”[41]. The output score range was between 0 and 1 with former being least cell penetrating and latter as most cell penetrating, We considered the score of more than 0.5 to be good and below that not-good for cell penetration for their analysis. Also, in house Python, R and Perl programmes were used in several components of the work like calculating the compositions, physicochemical properties or creation of the graphs [42].

### Organization, implementation and availability of the PEPsRNA web server

The main aim of this web server is to provide the information regarding the peptides used in the delivery of siRNAs. So this web server is mainly focused on the peptide based delivery and the siRNAs. Also, there were various other non-peptide molecules that were used in addition to the main peptides, which are also included (**Figure 1**).

We have organized the server sequentially with all the broad categories specified in the header. On the home page, the links of “Browse”, “Search”, and other analysis tools including “Help” page etc. provided [39]. The user can navigate through the options provided to browse, search, analyze and fetch the data.

MySQL supports the data in PEPsRNA webs server with sustenance from “LAMP (Linux, Apache, MySQL, PHP) software” package. It is presented by the help of front-end languages like HTML, CSS, JS, Java along with PHP, Python, R and Perl etc. working in synchrony with it. The data is available for educational and scientific use at the following web address: http://bioinfo.imtech.res.in/manojk/pepsirna.

## Supplementary data

Supplementary material:

## Acknowledgment

Supported by Council of Scientific & Industrial Research (CSIR) Institute of microbial technology (IMTECH) (OLP143), and Department of biotechnology (DBT) GAP0001

## Database URL

http://bioinfo.imtech.res.in/manojk/pepsirna

