## Supplemental File for "PEPsRNA: A computational resource of experimentally tested peptide based siRNA delivery"

**Supplementary Material**

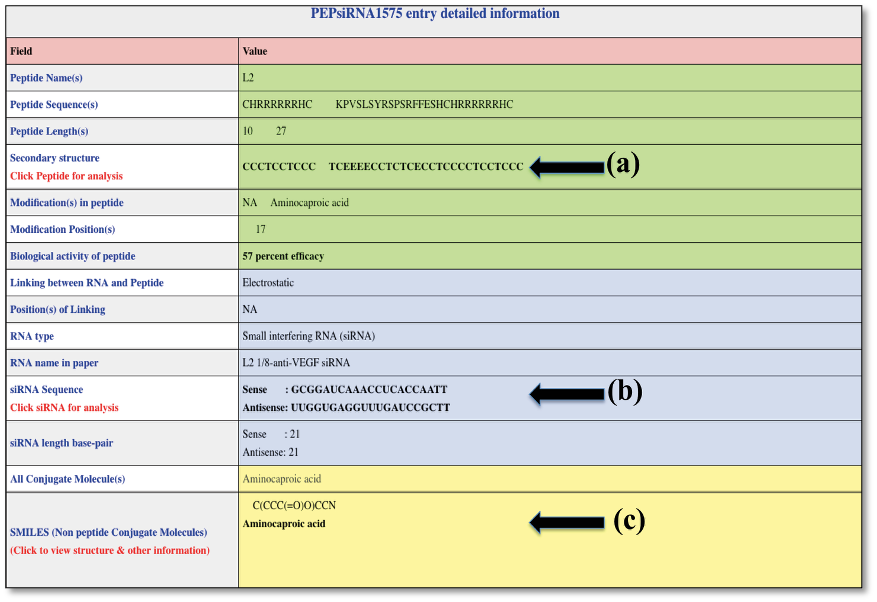

**Supplementary Figure S1.** The visual representation of the individual data entry with information about Peptide (a), siRNA (b) and non-peptide conjugate moiety (c) shown in different colors. On clicking the value in fields a, b and c (specified in red in figure above) their respective detailed information is displayed.

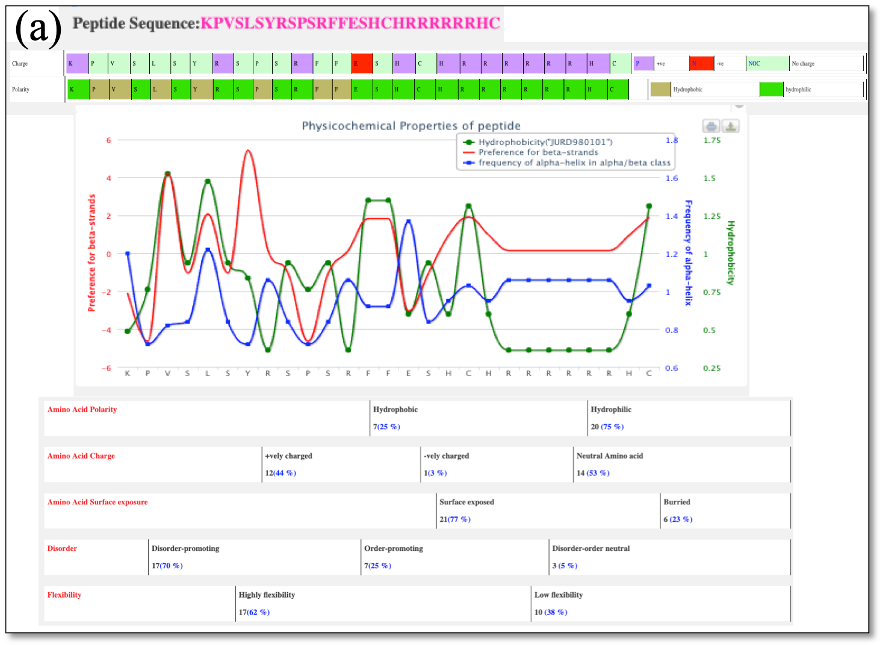

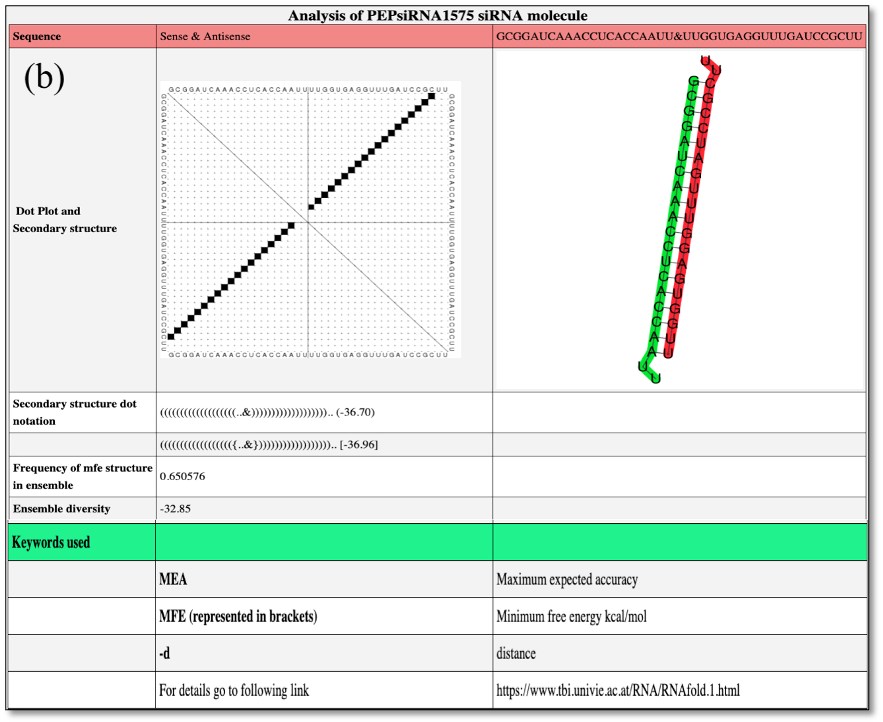

**Supplementary Figure S2.** Visual representation of the detailed analysis of the peptide and siRNAs on the web server; (All the data is not shown because of space limitation in page). (a) Peptide; with the sequence, its physicochemical properties in graphical and numerical values etc. (b) siRNA on the web server pages with its sequence, dot-plot, interaction of sense and antisense strand, secondary structure, minimum free energy etc.

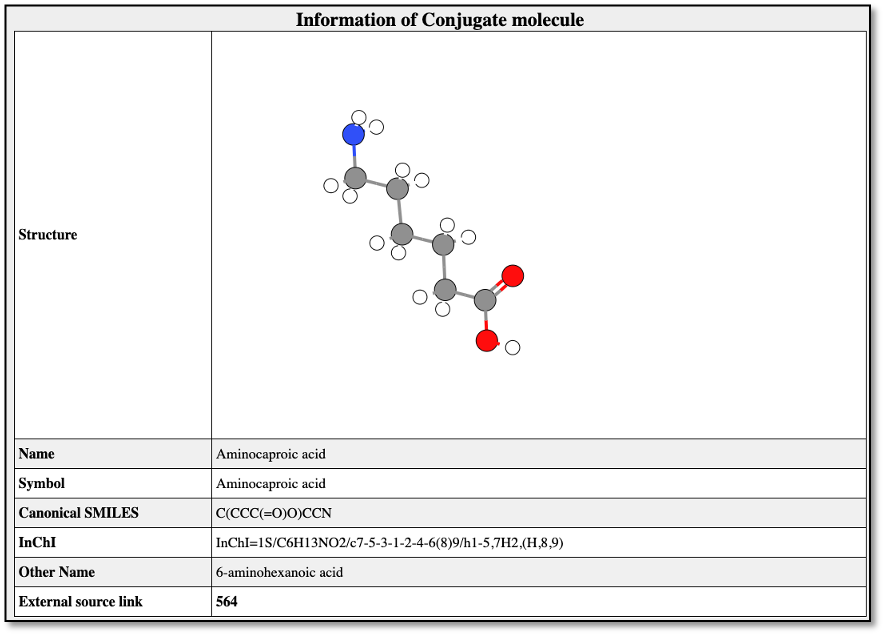

**Supplementary Figure S3.** Visual representation of the information of conjugate molecule other than peptide used in the delivery of siRNA.

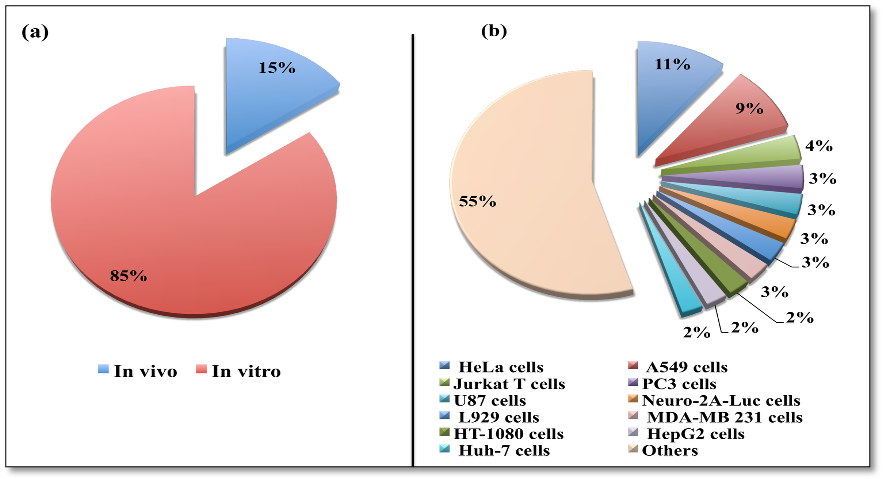

**Supplementary Figure S4.** The distribution of experiments for peptide based delivery as observed in this resource. (a) Distribution of *in vitro* and *in vivo* experiments and (b) Cell lines usage dispersal.

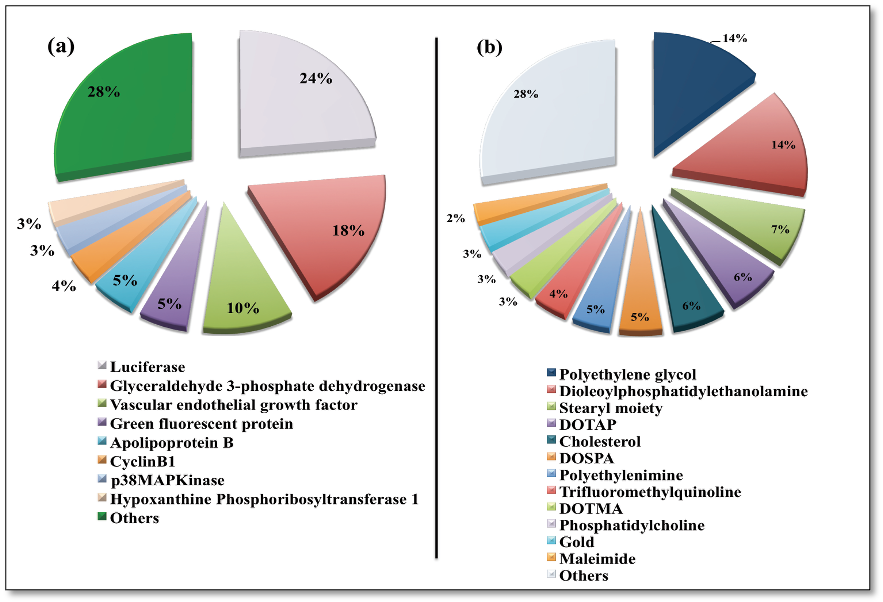

**Supplementary Figure S5.** Dispersal of (a) genes used for testing the delivery molecules and (b) the Conjugate molecules used in addition to the peptides for siRNA delivery.

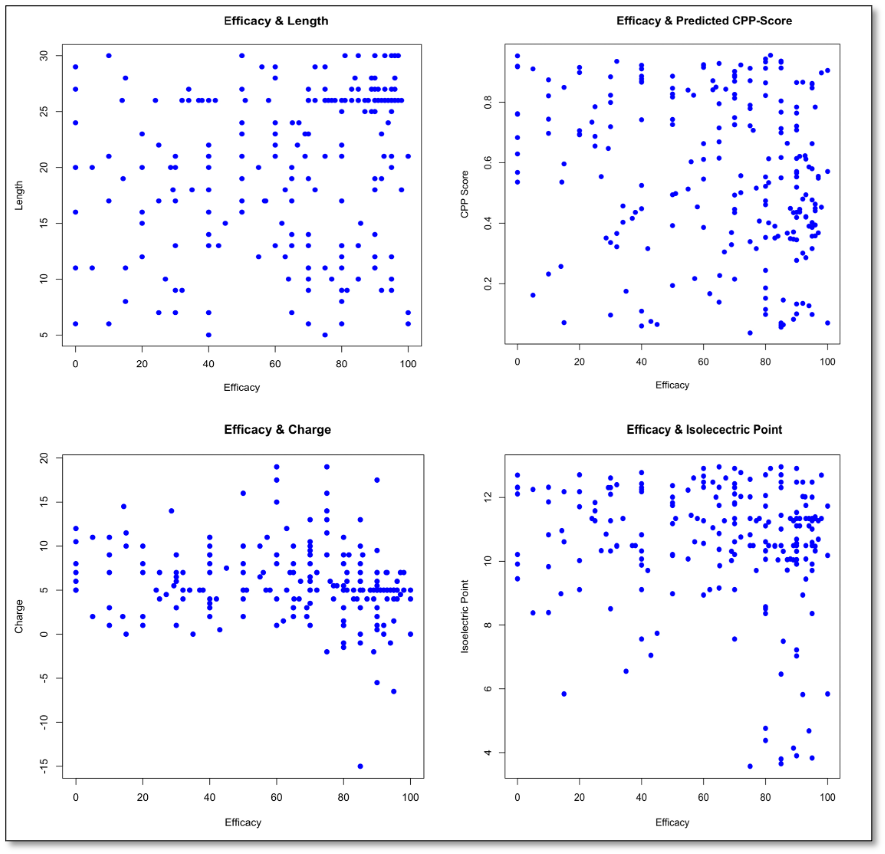

**Supplementary Figure S6.** Scatter plot analysis of the percentage peptide efficacy with respect to (a) Length (b) predicted cell penetrating score (CPP Score) (c) Charge (d) Isoelectric point (pI).

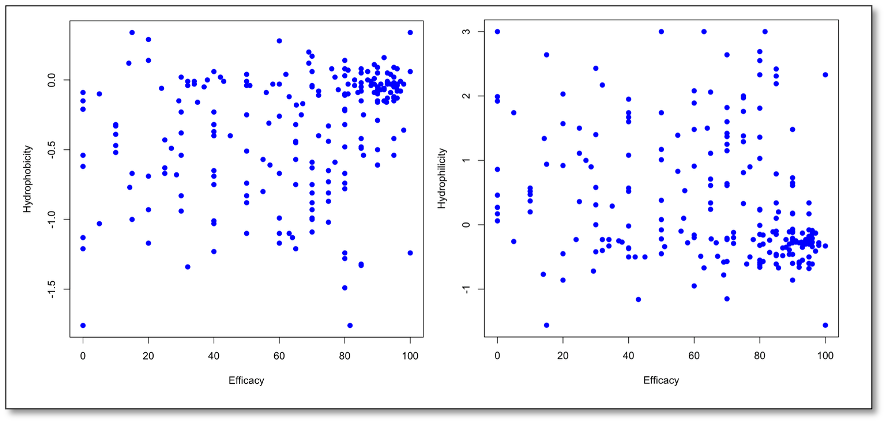

**Supplementary Figure S7.** Scatter plot analysis of peptides percentage efficacy in comparison to (a) Hydrophobicity and (b) Hydrophilicity.
